# Competition and the century-long decline of a once common lizard, *Sceloporus consobrinus*

**DOI:** 10.1101/2023.08.25.554877

**Authors:** Alexander H. Murray, Edita Folfas, Morgan A. Page, Zachary K. Lange, Joseph L. Mruzek, Luke O. Frishkoff

## Abstract

Habitat modification and climate change are primary mechanisms responsible for historical and ongoing population declines. Species interactions however, though difficult to study, may be of similar importance. Here we use a combination of historical species records, standardized transect surveys, and staged competition trials to assess the role of competition in recent population trends and distributions of two closely related lizard species: the prairie lizard (*Sceloporus consobrinus*) and the Texas Spiny lizard (*S. olivaceus*). Occurrence data reveals divergent population trends. *S. consobrinus* has decreased while *S. olivaceus* has increased in relative frequency over the last 100 years. We analyzed spatially-aggregated records of all lizards within the range of *S. consobrinus* to determine the role of climate suitability, climate change, landcover, and species interactions in shaping the occurrence patterns of *S. consobrinus*. In contrast to other lizard species, presence of *S. olivaceus* was associated with substantial reductions of *S. consobrinus* occurrence, and explains occurrence patterns better than either climate suitability or landcover. To test whether patterns of broad scale co-occurrence are indicative of local competitive exclusion we conducted 200m transects surveys to assess lizard communities and paired this with staged behavioral trials in the lab. Despite occurring in similar habitats, and across similar regions, transect surveys revealed lower-than-expected abundance of *S. consobrinus* on transects containing *S. olivaceus*, with both species co-occurring on only 2 of 176 transects. Shifts in habitat use implicate competitive displacement, with *S. consobrinus* occupying areas with three times less canopy cover at sites with *S. olivaceus* compared to those without. Finally, behavioral trials revealed competitive dominance of *S. olivaceus*, which controlled the prime basking position, and initiated more interactions that led to retreat or hiding by *S. consobrinus*. Our study implicates competitive interactions as in important force in structuring species’ distributions and population trends.

## Introduction

The last century has been marked by incredible changes in biodiversity writ large, emerging from shifts in community assemblages, population declines, changes in species distributions and species extinctions (Pereira *et al*., 2012; Dornelas *et al*., 2014; McGill *et al*., 2015; Dornelas *et al*., 2019). Explaining and ultimately predicting when and why some species decline is essential to both understand the ecology of the contemporary world and combat severe biodiversity loss. Habitat modification and climate change in particular are linked to many species’ declines (Pereira *et al*., 2012), yet substantial variation exists between species in how they respond to these global change drivers. While many species are “losers”, and suffer as a result of anthropogenic change, a few “winners” benefit and expand their population sizes and distributions (McKinney & Lockwood, 1999; Daily *et al*., 2001; Mendenhall *et al*., 2016). However, being a “winner” versus a “loser” is not necessarily a static trait. Across a given species’ range responses to the same type of anthropogenic induced change often varies (Frishkoff *et al*., 2015; Orme *et al*., 2019; Williams *et al*., 2021; Williams *et al*., 2022). Some of this intraspecific variation can be accounted for by proximity to physiological limits—for example deforestation, and the resulting local habitat warming due to increased solar warming, results in extirpation at the warm-edge of a species’ range, while the same deforestation may benefit the species towards the cold end (Frishkoff *et al*., 2015). Yet other cases of varying tolerance to global changes are not well understood, as changes in communities are often not explained by changing climate alone (Amburgey *et al*., 2018; Miller *et al*., 2018). Perhaps our inability to explain variation within species is hampered by our failure to consider the role that species interactions are playing, and how anthropogenic influence may be rewriting their strength and outcomes (Suttle *et al*., 2007; Blois *et al*., 2013; Alexander *et al*., 2016; Urban *et al*., 2016).

Climate plays a key role in determining species range limits (Angert & Schemske, 2005; Stanton-Geddes *et al*., 2012; Hargreaves *et al*., 2014; Lee-Yaw *et al*., 2016), community structure (Jackson *et al*., 2001), and species richness (Qian *et al*., 2007; Grace *et al*., 2016). In contrast, ecologists have vehemently disputed the role of competition in setting range limits, structuring communities and determining species richness (Connell, 1961; Wiens, 1977; Connell, 1980; Schoener, 1982; Connell, 1983; Roughgarden, 1983; Simberloff, 1983). Disagreement may stem in part from studies seeking to find an effect of competition but failing to do so, particularly in vertebrates (Tinkle, 1982; M’Closkey & Baia, 1987; Paterson *et al*., 2018). Failure to find strong competition amongst currently co-existing species should not be surprising, as species tend to minimize such interactions through niche partitioning. Such partitioning reduces the strength of competition between them, allowing them to persist in the presence of competitors (Connell, 1980). Evidence for such niche partitioning is widespread, and multidimensional, with species evolving or changing behavior to minimize overlap in diet (Huey *et al*., 1974), microhabitat use (Jenssen, 1973; Salzburg, 1984), thermal preferences (Watson & Gough, 2012) and morphology (Huey & Pianka, 1977). The tendency for competition to lead species to partition niches may make it difficult to detect the population effects of present-day competition, obscuring its importance in structuring communities and driving turnover across space. Further, the strength and impact of competitive impacts themselves may strengthen or weaken through time, especially during periods of environmental change and redistribution. Changes in abiotic conditions (Taniguchi & Nakano, 2000), habitat structure/heterogeneity (Petren & Case, 1998), and resource availability may all tip the scales, altering dominance between species pairs and drive once non-interacting species into competitive dominant and subordinate relationships. As human impacts across the globe continue to increase, we are altering the conditions under which species interact, making it likely that species interactions are changing in turn, yet we currently know little of their extent or consequences (Blois *et al*., 2013; Alexander *et al*., 2016).

We examine the role of competitive interactions in structuring communities and geographic distributions through time, using two closely related and ecologically similar lizard species: the prairie lizard (*Sceloporus consobrinus,* Baird and Girard 1854) and the Texas Spiny Lizard (*Sceloporus olivaceus,* Smith 1934). Lizards are often subjects for the study of competition (Pacala & Roughgarden, 1982), vary greatly in how they respond to land use change (Doherty *et al*., 2020), and can be highly sensitive to changing environmental conditions (Sinervo *et al*., 2010; Walker *et al*., 2015). Their sensitivity to anthropogenic change is likely amplified by their relatively low mobility, leaving them unable to rapidly track changing availability of resources in space, in contrast to groups like birds (Blake & Hoppes, 1986; Kasper Thorup, 2017). This limitation is crucial with regards to competition, because resource availability often determines lizard abundance and strength of competition (Dunham, 1980; Guyer, 1988). Anecdotes, personal communications, and isolated reports all suggest that *S. olivaceus* abundance has been increasing, while *S. consobrinus* has been declining (Mora, 1991). Some of this trend is likely due to land use change—*S. olivaceus* appears more tolerant of urban areas than *S. consobrinus* (personal observation). However, even in localities with no history of land-use change, *S. consobrinus* has disappeared, and *S. olivaceus* remains (Mora, 1991). This anecdotal pattern conforms to expectations if interspecific competition were responsible. Such an interpretation is further supported by the size and shape of these species’ ranges. *S. consobrinus* is widespread between the Mississippi river and the Rocky mountains. However, records suggest much lower occurrence frequency or even absence in an internal portion of their range in central Texas—a geographic region occupied by *S. olivaceus*, whose range runs in a narrow east-west band from Central Texas south into Mexico.

What is therefore missing is (i) a formal analysis verifying these broad-scale patterns through time and over space, (ii) standardized local surveys to indicate lack of co-occurrence between species at the scale of individual lizard home ranges along with evidence of habitat-based niche partitioning that could come about due to competition, and (iii) evidence that behavioral interference competition between these species actually occurs, and that *S. consobrinus* in the inferior competitor. While as in any historical science, these forms of correlative evidence cannot prove that competitive interactions *caused* the decline of *S. consobrinus*, if all three lines of evidence point in the same direction it would build a strong case implicating competition as a likely cause.

Here, we combine occurrence data, transect surveys and behavioral trials to work from macro- to micro-scale, testing our core hypothesis that *S. consobrinus* and *S. olivaceus* compete and that such competition is structuring range limits and contributing to the decline of *S. consobrinus*. First, we use occurrence data to test our hypothesis that relative rates of occurrence of *S. consobrinus* are decreasing through time (and *S. olivaceus* is increasing) to confirm anecdotal reports. Second, we use these records to assess broad-scale co-occurrence, asking whether *S. consobrinus* occurrence is lower than expected in areas which are otherwise suitable for it if *S. olivaceus* is also present. Third, we use standardized transect surveys to test whether broad-scale species co-occurrences are representative of fine-scale community structure, and test our hypothesis that abundance of *S. consobrinus* will be lower on transects which contain *S. olivaceus*. Finally, we use behavioral trials to test whether *S. olivaceus* and *S. consobrinus* indeed compete directly via behavioral interference—and specifically whether *S. olivaceus* is the superior competitor.

## Methods

### Study system

We center our study across the state of Texas where the geographic ranges of *S. consobrinus* and *S. olivaceus* overlap. Texas spans a gradient in precipitation, with related variation in habitat across the state and corresponding turnover of lizard communities. In the last century Texas has experienced substantial change, from major human population increases and resulting urban sprawl, shifts in precipitation patterns, and the introduction of non-native species.

*Sceloporus consobrinus* is generally regarded as common throughout Texas, in habitats ranging from xeric scrub, to prairies, to broadleaf and pine forests. In conjunction with its wide distribution, *S. consobrinus* exhibits incredible beauty and phenotypic diversity. They are small (max SVL= 68mm; (Lawrence LC Jones & Lovich, 2009), “semi-arboreal” lizards. They can be highly arboreal in some parts of their range but in more sparsely vegetated areas are more terrestrial. They belong to the *Sceloporus undulatus* group (Leache, 2009), which is distributed throughout the US and in most places contains a single small to medium sized lizard ecologically similar to *S. consobrinus*. In central Texas, *S. consobrinus’*s range overlaps with the much larger and closely related species, *Sceloporus olivaceus* (max SVL= 124mm; (Kennedy, 1973), which is endemic to wooded habitats of central Texas and Northern Mexico (Figure 1). Both species are diurnal insectivores, and in Texas can be active year-round, though with less activity in the cooler winter months. Little is known about how territorial or aggressive these species are, but they likely defend territories, similar to other *Sceloporus* species (Ruby, 1978). The fine-scale distribution of *S. olivaceus* individuals suggests this to be the case, as single trees tend not to possess multiple males (Blair, 1960).

**Figure 1.**
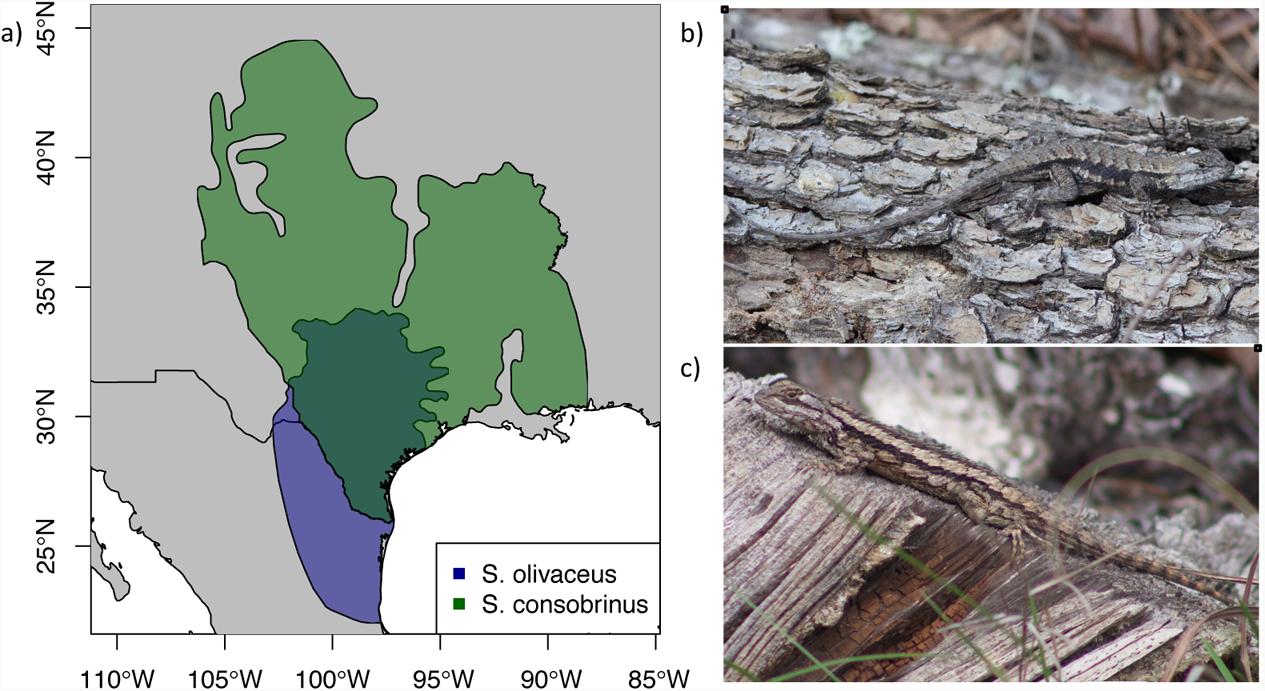
a) Distributions of *Sceloporus olivaceus* and *Sceloporus consobrinus.* b) Male Prairie lizard (*Sceloporus consobrinus*), Jasper county, TX. c) Male Texas Spiny Lizard (*Sceloporus olivaceus*), Travis county, TX.

### Trends in occurrence

#### Data

In order to document divergent trends in occurrence during the 20^th^ and 21^st^ centuries we assessed time- and geo-referenced occurrence data. First, we assessed whether the general trends in records of *Sceloporus consobrinus* are decreasing through time and whether the trends in records for *S. consobrinus* differ from *S. olivaceus*. To do so we downloaded occurrence data from GBIF for all lizards on December 3^rd^, 2021. We constrained the area of study to the distribution of *S. consobrinus* and dropped records which fell outside of this distribution, as well as records which lacked coordinates. We retained 91,660 records of lizards between 1920 and 2021. Records of *S. undulatus* falling within the distribution of *S. consobrinus* were reclassified as *S. consobrinus* to reflect updated taxonomy (Figure S1). To assess the trends in observations over time we grouped the data into ten-year periods from 1920-2021, except for one two-year period, 2020-2021. The time period with the fewest records was 1920-1929 with 1,902 records and 2010-2019 had the most with 29,499 records.

#### Analysis

We gauged potential changes in population trajectory of our focal species based on changes in relative abundance versus all lizards in the study area. We used a linear model, with a Gaussian error distribution and the *glm* function (R Core Team, 2016). We used log of relative abundance (i.e. log(focal species observations / total lizard observations) as our response variable, and included an interaction with year and species (i.e. *S. consobrinus* or *S. olivaceus*). These and all subsequent analyses were performed in R version 3.6.2.

### Current Distribution

#### Pseudo-sites

To understand what explains the current distribution of *Sceloporus consobrinus*, we spatially and temporally aggregated records into pseudo-sites, allowing us to assess the influence of land-use, climate and species interactions on the current distribution of *S. consobrinus*. In essence, pseudo-sites are groupings of observations in the same general geographic area, that are used to control for sampling effort, thereby ensuring that the absence of *S. consobrinus* was not simply due to a lack of sampling for lizards in the general vicinity. To spatially aggregate our data we generated pseudo-sites by creating 100,000 random points within the distribution of *S. consobrinus* using the *randomPoints* function (Hijmans *et al*., 2017). We buffered our points by a 2km radius to allow for the grouping of near-by records. We further cleaned the GBIF dataset by dropping all records with a coordinate uncertainty greater than 2km and restricted the time period to only the most recent records, 2020-2021. For the two focal species, geographic outlier records were assessed visually to confirm ID, and records with incorrect ID were dropped or changed. We then intersected the occurrence records with the buffered pseudo-sites using the *intersect* function (Hijmans RJ, 2015). For each pseudo-site, we summed observations and dropped pseudo-sites containing fewer than 5 records. To ensure each pseudo-site represented a unique community, with no duplicate records, we spatially thinned out the data set by 4km using the *thin* function (Aiello-Lammens *et al*., 2015), we retained a total of 396 pseudo-sites.

#### Climate Suitability and Land-use data

We used Maxent (Phillips *et al*., 2006) to predict the climatic suitability for *S. consobrinus* and *S. olivaceus*. We cleaned the initial GBIF dataset by removing all records with >1km coordinate uncertainty, and geographic outliers, and correcting records to reflect current taxonomy. After thinning (1km), we retained a total of 2,527 records (Figure S3). Similar cleaning for *S. olivaceus* resulted in a total of 2,817 records (Figure S4). Climate data were obtained at 1km resolution from WorldClim (Fick & Hijmans, 2017). We selected climate variables based off stepwise exclusion using *vifstep* function from the package *usdm* (Naimi *et al*., 2014). The resulting final variable set for both species included: bio2, bio4, bio5, bio8, bio9, bio15, bio18. Calibration area was determined based off a minimum convex polygon which includes all occurrence points and a 200km buffer around the area. Occurrence records were split into 5 groups using a *kfold* function from the *dismo* package (Hijmans *et al*., 2017), and 80% of records were used for model training, 20% was used to evaluate the models. Models were run using the *maxent* function, output was formatted as a logistic output as it represents an attempt to determine probability that a species is in a location given the environmental conditions (Elith *et al*., 2011). We evaluated model performance with 10,000 background points and 20% of occurrence records for each species using the *evaluate* function (Hijmans *et al*., 2017). Values for AUC value for *S. consobrinus* was 0.749 (Figure S5), whereas the AUC for *S. olivaceus* was 0.909 (Figure S6).

We obtained land-use data for urban area, pasture, crop and forest at ~1km × 1km resolution for the year 2020 (Li et al 2022). Maxent predicted habitat suitability and land-use data were extracted to the pseudo-sites. For each time period we extracted mean annual temperature (MAT) and mean annual precipitation (MAP) to the sites. Historical climate data were obtained at 1km resolution (PRISM Climate Group, 2011). Changing climatic conditions may favor one species over another, and so we accounted for recent climate change by creating rasters representing changes in MAP and MAT by subtracting the mean for 1920-1929 from the mean value for 2010-2019 and extracted these values to our pseudo-sites.

#### Analysis

Generalized linear models were used to assess the factors responsible for the current distribution of *S. consobrinus*. We used presence/absence of *S. consobrinus* in the 396 pseudo-sites as the response variable, and evaluated whether (1) climate suitability, (2) degree of MAT and MAP change over the last century (3) land-cover type (4) presence/absence of 11 other lizard species, influence the presence and absence of *S. consobrinus*. In all models we included number of records at the site to control for the influence of sample depth. We evaluated model performance based of off AIC values, and calculated pseudo-R^2^ values through the *pR2* function (Jackman, 2015).

### Mechanisms driving trends

Pseudo-sites were generated and cleaned as above for the current distribution analysis, but using data from all years between 1920-2021, grouped into ten year time periods. We retained a total of 1340 pseudo-site year combinations between 1920-2021 (Figure S2). The average number of records per site was 16 and the range was 5 to 450.

#### Analysis

To test if occurrence of *S. consobrinus* is changing through time we used linear models with a binomial error distribution and presence/absence of *S. consobrinus* in the 1340 pseudo-sites as the response variable. First, we tested whether occupancy of *S. consobrinus* in pseudo-sites has decreased through time while controlling for the influences of habitat suitability, sampling and landcover as predictors of the occurrence of *S. consobrinus*. Next, we assessed whether the effect size of *S. olivaceus* on *S. consobrinus* has changed through time, through including an interaction between *S. olivaceus* presence and time in the model. Landcover-type may influence the competitive outcomes between species. We therefore tested whether changing landcover is contributing to changing trends between *S. olivaceus* and *S. consobrinus*, including an interaction between *S. olivaceus* presence and amount of urbanization. This assesses whether urban areas have a more negative impact on *S. consobrinus* when *S. olivaceus* is present. Finally, we tested whether climate conditions altered the rate of exclusion by *S. olivaceus*, including interactions with climate and *S. olivaceus* presence, to determine if co-occurrence is more likely in some climates compared to others. We evaluated model performance based of off AIC values. Final models included climate suitability for *S. consobrinus, S. olivaceus* presence, land-cover, as well as interactions between urbanization and *S. olivaceus* presence, and interactions between climate and *S. olivaceus* presence.

### Lizard Community Surveys

#### Transect

Occurrence data can offer a glimpse into communities at coarse geographic and time scales yet lacks the ability to display the nuance of habitat partitioning within communities at finer scales. To better understand the associations between species, their microhabitat use, and partitioning within the landscape at the spatial and temporal scales over which individual lizards actually interact with one another, we used transect surveys between March 2020 and June 2022.

Transects consisted of 200m surveys, broken into four, 50m sub-transects. Sub-transects occurred in consistent habitat and allowed finer resolution data on species co-occurrence. Individual lizard surveys were conducted by a single individual walking slowly and scanning all substrates within line of sight. All cover objects of suitable sizes (>500cm^2^) within 2m of the transect line were flipped to check for lizards. When a lizard was observed, it was photographed and microhabitat data was taken (substrate, perch height, perch diameter, and perch temperature). Surveys averaged ~120 minutes and were only performed in habitat representative of the natural vegetation in a region. A total of 37 sites (with 1-36 transects per site) were visited for surveys spanning most of the range of *S. consobrinus* in Texas. In locations with extensive natural habitat we conducted many surveys, with multiple types of habitats sampled (canyons, riparian forest, prairie etc.), to account for all natural habitat types at the site. To maximize detection probability on transects, surveys were only performed during the months of the year when our focal species are most active (March-October), and under suitable weather conditions (ambient temperature > 20 degrees C°, and no rain). A total of 176 surveys are included here.

We recorded vegetation data to understand how vegetative structure influences the lizard community, measurements were taken at the midpoint of each sub-transect. We recorded canopy height of the tallest tree (using a TruPulse Hypsometer) and canopy cover (using a convex Spherical Crown Densiometer). Canopy cover was taken with four measurements in each subtransect, one facing each cardinal direction.

#### Analysis

Data were analyzed using generalized additive mixed models using the *gamm* function (Wood, 2015) and a Poisson distribution to determine the influence of *Sceloporus olivaceus* on the abundance of *S. consobrinus*, while controlling for the canopy cover, climatic suitability, and location. We accounted for the influence of climate on species abundance by including MAT and MAP of the transect in the model. All transects were in natural vegetation, but we controlled for the effect of landscape level urbanization by extracting the amount of urbanization within 2km surrounding our transects. To test if at local scales *S. consobrinus* are being excluded from otherwise suitable habitat we included presence/absence of *S. olivaceus* at the transect level as a predictor variable. In addition to the influence of *S. olivaceus* on *S. consobrinus* we tested for species associations with the other 10 most common species.

We also compared habitat use between *S. olivaceus* and *S. consobrinus* to determine if they have shared preference based off perch height, perch temperatures and canopy cover on transects where they are found. To do this we used t-tests to compare the means between species, treating individual observations of lizards as samples. Since competition can alter habitat use, we tested whether habitat use of *S. consobrinus* differs in areas which lack *S. olivaceus* compared to those where it is present. To do this we grouped observations of *S. consobrinus* into sites which contain *S.olivaceus* (those where we observed *S. olivaceus*) and those where *S. olivaceus* is absent and compared canopy cover between the two.

### Competition Trials

We conducted competition trials between *S. olivaceus* and *S. consobrinus* to assess which species is dominant based off rates of aggression and retreats, and changes in habitat use when in the presence of individuals of a competitor versus when alone.

#### Data

Competition trials took place inside of a 100×45×45cm enclosure containing a variety of substrates, including leaf litter, dirt, rock, and branches of 3 different sizes (Figure S7). Each enclosure had a heat lamp placed directly on top of the cage as *Sceloporus* species have high thermal needs compared to the ambient temperature. Trials lasted 120 minutes and were recorded using a set of security cameras so lizard habitat-use and interactions could be scored later. Lizards were only used in one trial per treatment. A total of 21 trials were performed with adult male *S. olivaceus* paired with adult male *S. consobrinus*, 9 trials were run with single *S. consobrinus* on their own.

#### Analysis

Competition trials were scored in 1-minute increments every 10 minutes, starting with the first minute of the trial. During each scoring period we recorded the substrate that the lizard was on, and noted any interactions between individuals. Interactions were recorded to include which lizard initiated the interaction, what the initial behavior was, and how the other lizard responded. Behaviors by lizards initiating interactions were given numerical values based off aggression level: 1 (approaching the other lizard, pushups, and headbobs), 2 (approaching while head bobbing or pushuping, or approaching and touching), 3 (biting). Responses to the initial behavior was recorded as: 0 (no response), −1 (retreat but maintain presence on same substrate), −2 (retreat and change substrate), −3 (retreat and hide), or positive values as above if the response was a counter-aggressive display. We tested the hypothesis that *S. olivaceus* is the dominant competitor by using t-tests to compare mean levels of aggression, response to aggression within trials, and habitat use withing trials. T-tests were also used to determine if the substrate use differed in two species trials versus single species trials.

## Results

### Trends in occurrence and Distribution of *S. consobrinus*

As a proportion of all lizards, *Sceloporus consobrinus* has declined in frequency of observations over the last century (p-value < 0.001, Figure 2a). It made up more than 20% of lizard records in the early decades of the 20^th^ century, but only ~ 5% in the 2010s. Over this same time *S. olivaceus* has increased from less than 5% to more than 10% of lizard records. Changes in landcover, climate, or species interactions may all potentially explain the decline of *S. consobrinus.* We first examined how each of these classes of predictors explained the current distribution of occurrences. Surprisingly, climate alone does a very poor job of explaining *S. consobrinus* presence at the 396, 2km-radius “pseudo-sites” (R^2^ = 0.008; p >0.05, Table 1). Urbanization, in contrast, has a strong negative influence on *S. consobrinus* (p-value <0.001, R^2^= 0.14), though other forms of land use have little influence (Table 1). While present day climate suitability had limited predictive capacity, the degree of climate change over the last century was a strong predictor of their current distribution. Areas which had warmed recently (p-value<0.001, R^2^ = 0.11), and those which had gotten wetter recently (p-value<0.001, R^2^ = 0.08) were less likely to be occupied by *S. consobrinus* (Table 1).

**Figure 2.**
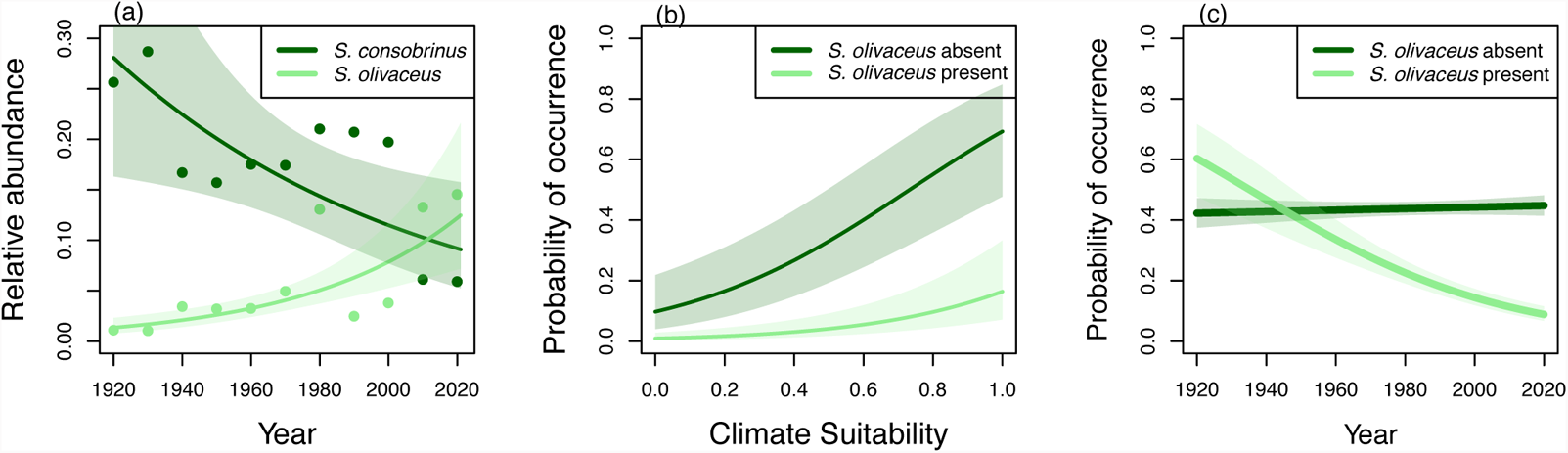
a) Trends in records of *Sceloporus consobrinus* and *S. olivaceus* over the last 100 years relative to all lizard records within the distribution of *S. consobrinus*. Lines represent model-based predictions of relative abundance for each species based off decadal values of relative frequency, shaded areas represent 95% confidence intervals from model predictions. Points are the raw values for relative frequency of each species for each ten-year period. b) Model-based predictions of the probability of occurrence of *S. consobrinus* in areas without urbanization based off Maxent predicted values for climate suitability (range in pseudo-sites 0.12-0.92, mean=0.61) while controlling for effects of sampling effort. Shaded areas represent standard error. c) Change in strength of competitive exclusion of *S. consobrinus* by *S. olivaceus* in natural areas(urban=0) through time, while controlling for the effect of climate suitability, and sampling.

**Table 1.** Results from models testing the current distribution of *S. consobrinus*, 2020-2021. Number of pseudo-sites=396. *P-value <0.05 *, <0.01**, <0.001\*\*\**.

| Model # | Type | AIC | R2 | ΔAIC | 1 | 2 | 3 | 4 | 5 | 6 | 7 |
| --- | --- | --- | --- | --- | --- | --- | --- | --- | --- | --- | --- |
| 1 | climate suitability | 271.5 | 0.008 | 86.1 | hbs | sampling |  |  |  |  |  |
| 2 | climate suitability +landcover | 238.5 | 0.14 | 53.1 | hbs | sampling | -urban*** |  |  |  |  |
| 3 | climate suitability+ climate change | 246.9 | 0.11 | 61.5 | hbs | sampling | -MAT change*** |  |  |  |  |
| 4 |  | 255.4 | 0.08 | 70 | hbs | sampling | -MAP change*** |  |  |  |  |
| 5 |  | 229.6 | 0.18 | 44.2 | hbs | sampling | -MAT change*** | -MAP change *** |  |  |  |
| 6 | climate suitability+landcover+ climate change | 221.9 | 0.22 | 36.5 | hbs | sampling | -urban** | -MAT change*** | -MAP change** |  |  |
| 7 | climate suitability+ landcover+ species associations | 212.4 | 0.24 | 27 | +hbs** | sampling | -urban*** | -S. olivaceus *** |  |  |  |
| 8 |  | 221.6 | 0.21 | 36.2 | hbs | sampling | -urban** | -A. carolinensis *** |  |  |  |
| 9 |  | 236.2 | 0.16 | 50.8 | hbs | sampling | -urban*** | +P. fasciatus* |  |  |  |
| 10 |  | 236.1 | 0.16 | 50.7 | hbs | sampling | -urban*** | +Scincella lateralis* |  |  |  |
| 11 |  | 194.8 | 0.33 | 9.4 | +hbs* | sampling | -urban** | -S. olivaceus *** | -A. carolinensis *** | +Scincella lateralis* |  |
| 12 | climaye suitability + species associations | 236.8 | 0.15 | 51.4 | +hbs** | sampling | -S. olivaceus *** |  |  |  |  |
| 13 | climate suitability + climates suitability of S. olivaceus | 217 | 0.22 | 31.6 | +hbs*** | sampling | -hbs S. olivaceus *** |  |  |  |  |
| 14 | climate suitability+landcover+climate change+species associations | 193.5 | 0.34 | 8.1 | +hbs* | sampling | -urban** | -S. olivaceus *** | -A. carolinensis *** | Scincella lateralis | MAT ct |
| 15 | climate suitability + climates suitability S. olivaceus +landcover+ species interactions +climate change | 185.4 | 0.37 | 0 | +hbs* | sampling | -hbs S. olivaceus *** | -urban** | -A. carolinensis ** | Scincella lateralis | MAT ct |

On top of effects of climate and landcover, species interactions may also limit species distributions. We therefore tested for associations between species within pseudo-sites, while controlling for climate suitability, sampling amount, and urbanization in all assessments. Of the 11 common Texas lizard species analyzed, four had significant associations with *S. consobrinus* presence. The strongest correlation was with *S. olivaceus* (p-value <0.001, R^2^=0.24) for which occurrence of *S. olivaceus* was associated with diminished occurrence of *S. consobrinus*. Green anoles (*Anolis carolinensis)* also had a strong negative correlation in occurrence with *S. consobrinus* (p-value<0.001, R^2^=0.21). In contrast, both the little brown skink (*Scincella lateralis*, p-value<0.05, R^2^= 0.16) and the five lined skink (*Plestiodon fasciatus;* p-value<0.05, R^2^= 0.16) had positive association with *S. consobrinus*. Including multiple species within one model suggested a strong negative effects of *S. olivaceus* and *A. carolinensis*, and a weak positive influence of *S. lateralis*, R^2^=0.33 (Table 1). Our best model included landcover, species associations and climate change, R^2^= 0.34, though climate change no longer had a significant influence on *S. consobrinus* occurrence once species associations were incorporated (Table 1).

### Potential mechanisms driving declines

Having observed a decline in *S. consobrinus* through time (p-value<0.001, R^2^=0.09; Table 2), we sought to understand which factors could explain declining trends. The average amount of urban cover for sites in 1920 (6% urban) was far less than today (56% urban) suggesting that urbanization may be in part responsible for the decline of *S. consobrinus*.

**Table 2.**
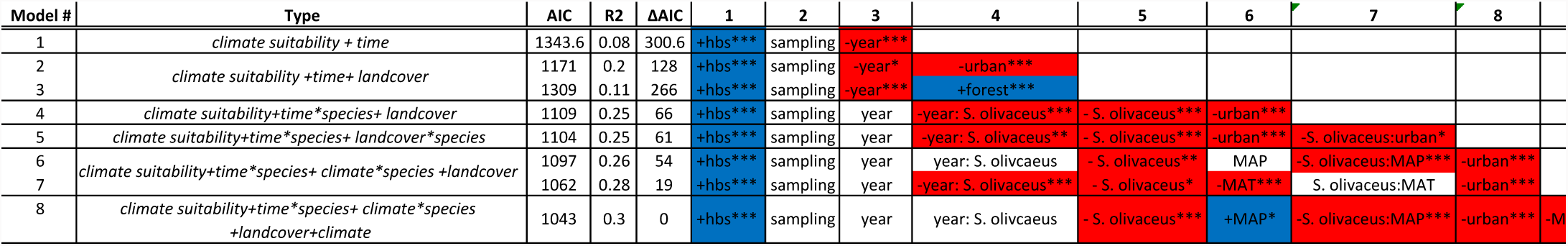
Results from models testing the occurrence of *S. consobrinus* through time, 1920-2021. Number of pseudo-sites=1340. *P-value <0.05 *, <0.01**, <0.001\*\*\**.

Likewise, relative records of *S. consobrinus* versus *S. olivaceus* as a fraction of total lizard records move in opposite directions, with *S. olivaceus* increasing over the last 100 years, as *S. consobrinus* declines (interaction between year and species identity predicting relative frequency, R^2^=0.57, p-value <0.001, Figure 2a)—as would be expected if *S. consobrinus* were being impacted by a competitor. Interestingly, when examining these historical records, instances of co-occurrence between *S. consobrinus* and *S. olivaceus* in 2km radius “pseudo sites” were more common during the early 20^th^ century than they are today (Interaction effect between year and *S. olivaceus* presence on predicting occurrence of *S. consobrinus*, p-value <0.001, R^2^=0.25, Figure c).

One possibility is that changing landcover and climate over the last century have been favorable for *S. olivaceus* allowing it to exclude *S. consobrinus* in areas that have experienced changes. We tested whether urbanization impacts the frequency of co-occurrence, and we found that in urban areas the effect of *S. olivaceus* is amplified, further decreasing the already low capacity for *S. consobrinus* to occur (p-value<0.001, model R^2^= 0.25, Figure 3). We further tested for interactions between climate and *S. olivaceus* occurrence to determine if certain climatic conditions allow for co-occurrence while others do not. While warmer areas were less likely to possess *S. consobrinus* (Figure 4), we detected no interaction between *S. olivaceus* presence and mean annual temperature (p-value > 0.05; Table 2). Wetter areas were more likely to possess *S. consobrinus*, but wetter areas where *S. olivaceus* occurred were highly unlikely to contain *S. consobrinus* (*S. olivaceus*-by-MAP interaction effect: p-value<0.001, R^2^=0.30) (Table 2, Figure 4). These results suggest that both urbanization and high precipitation may enhance the negative effect of *S. olivaceus* on *S. consobrinus* occurrence. While negative co-occurrence of observation records at broad spatial scales seems to implicate competition between *S. olivaceus* and *S. consobrinus*, proper evaluation of this hypothesis requires standardized surveys to evaluate co-occurrence at fine scales.

**Figure 3.**
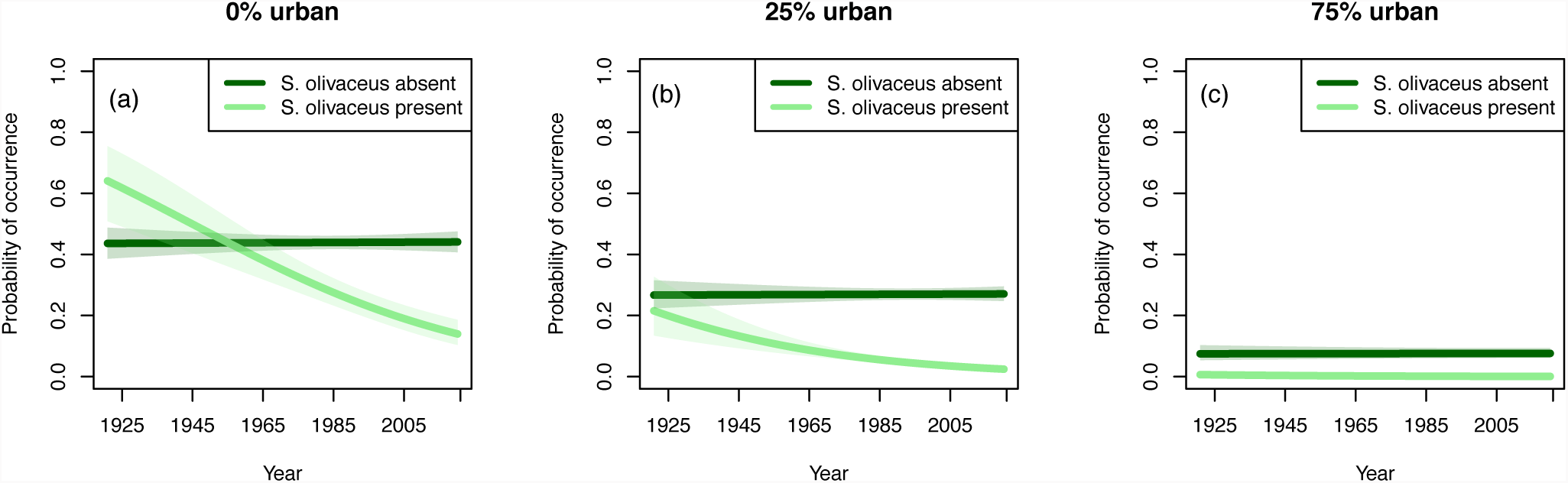
a) Predicted probability of *S. consobrinus* occurrence over time in areas that have no urbanization based off presence or absence of *S. olivaceus* while controlling for effects of climate suitability and sampling. Shaded areas represent standard error. b) Predictions made for areas which are 25% urban. c) Predictions for areas where 75% is urban. Pseudo-sites ranged from 0-100% urban, with a median of 9.8%, 25% of pseudo-sites were below 1% while 75% of the pseudo-sites were above 76%.

**Figure 4.**
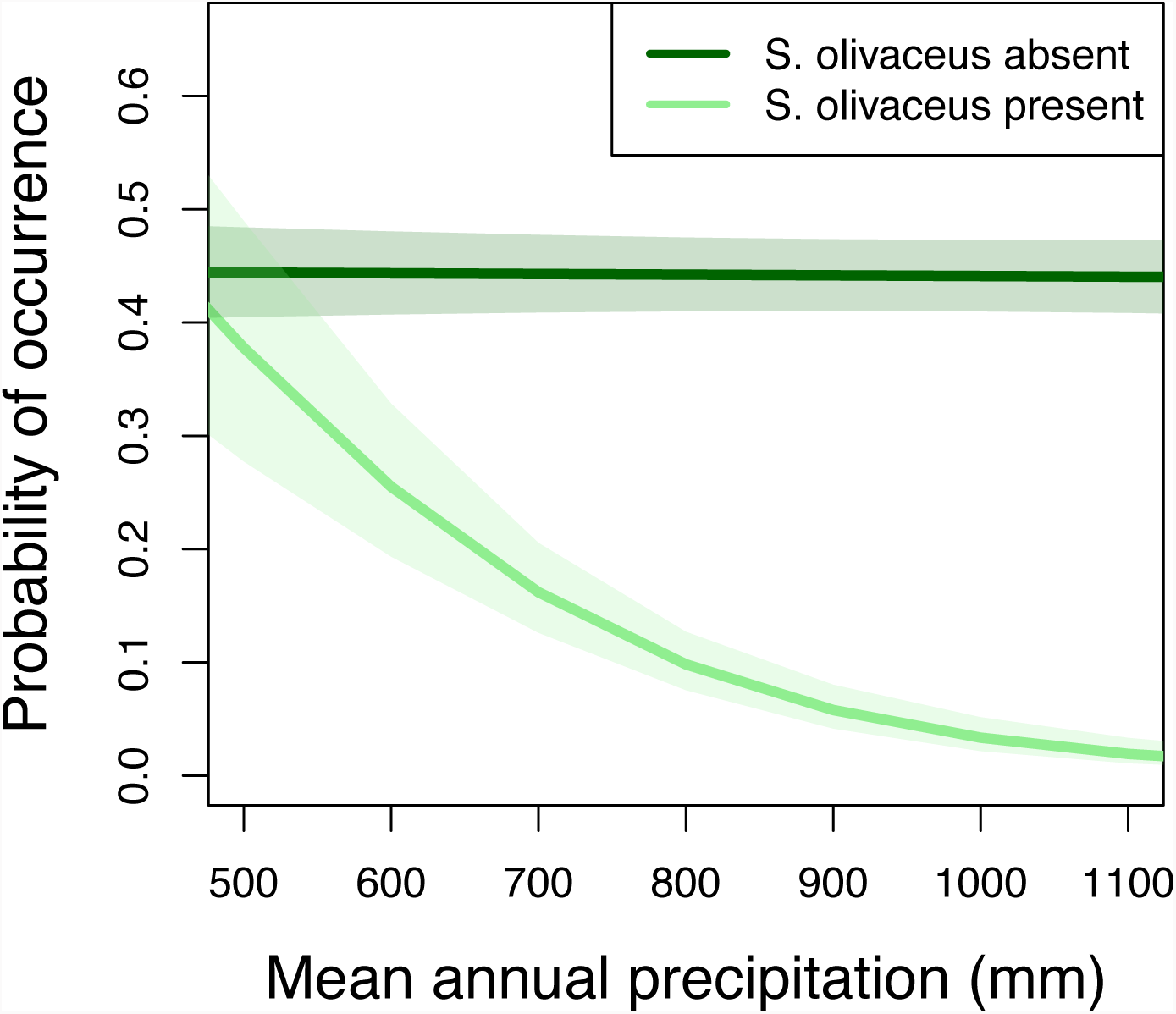
Model-based predictions of the probability of occurrence of *S. consobrinus* in areas without urbanization based off presence or absence of *S. olivaceus* and mean annual precipitation (range confined to the 5% and 95% for MAP at pseudo-sites where *S. olivaceus* is present). Shaded areas represent standard error.

### Lizard Communities

We analyzed data from 176 transect surveys to assess if the trends of exclusion of *S. consobrinus* by *S. olivaceus* is supported by patterns of co-occurrence at smaller spatial scales, in the same 200m transects during the same day (Figure 5). We tested if microhabitat use between the two species was different based off perch height and substrate temperatures of individuals observed on transects. Average perch height was not significantly different (*S. consobrinus* mean = 107cm, 25-75% = 30-118 cm, *S. olivaceus* mean = 135cm, 25-75%=50-180 cm, p-value= 0.12) and overlapped substantially. Likewise, both species were most frequently observed on woody substrates, often on tree trunks, fallen logs or branches. (Figure 6a). Substrate temperatures of field active individuals were also not significantly different between the two species (*S. consobrinus* mean = 33.7, 25-75% = 30.9-35.5 °C, *S. olivaceus* mean = 33.0, 25-75%= 31.6-34.9 °C, p-value=0.18). Based on our data of microhabitat use, niche similarity between *S. consobrinus* and *S. olivaceus* is greater than between any other common co-occurring species in the Texas lizard fauna (Figure 6a). In contrast, *A. carolinensis,* which was implicated as a potential competitor in broadscale analyses above, has substantially different thermal preferences, even though structural habitat use is not dissimilar.

**Figure 5.**
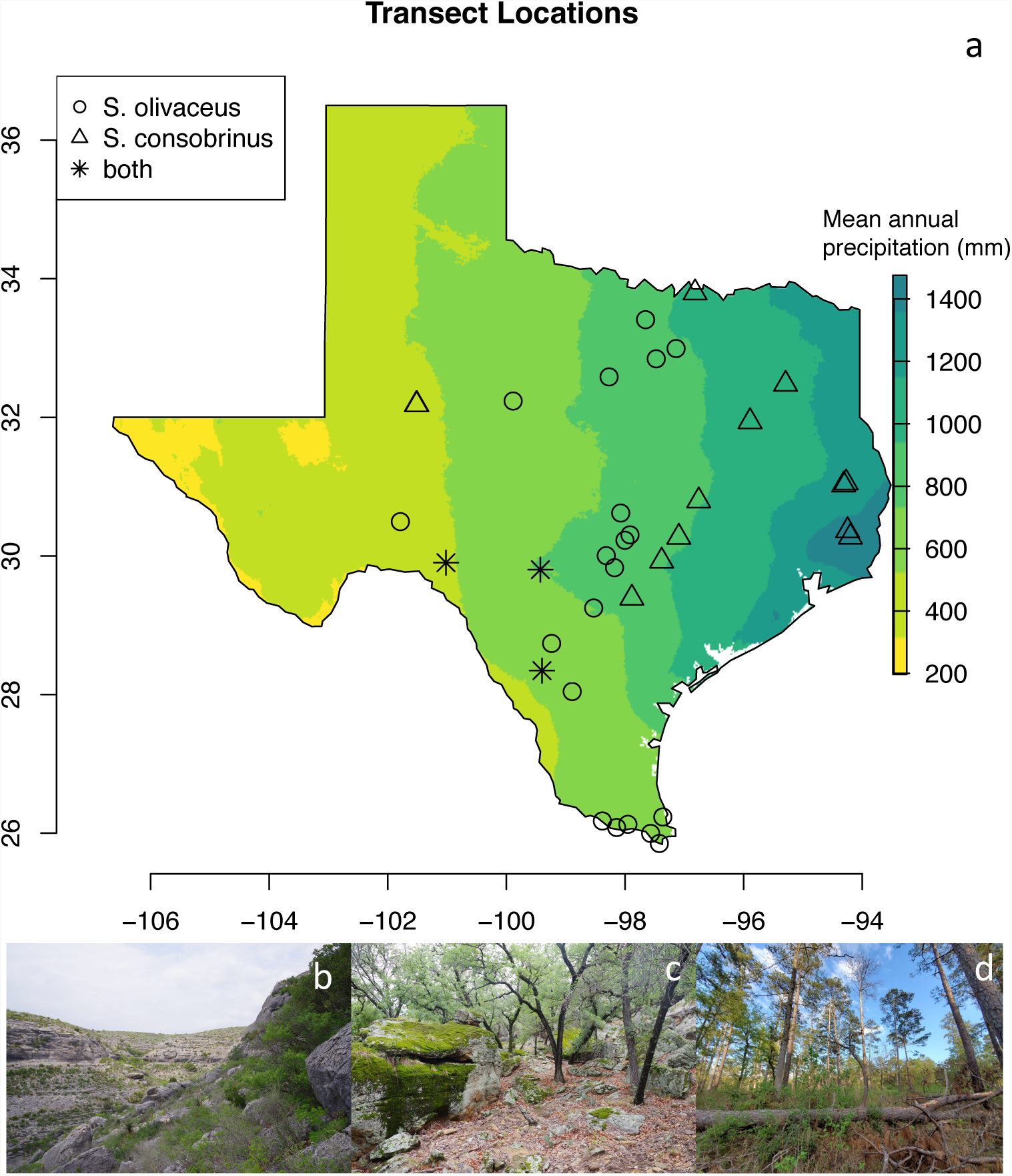
a). Distribution of sites where either *S. olivaceus* or *S. consobrinus* was observed. b). The Dolan Falls Preserve, habitat representative of the canyons in the western portion of the Edwards Plateau, forested base of canyon contains *S. olivaceus* open plateaus on top have *S. consobrinus*. c) Roger Fawcett WMA, central Texas cross timbers, habitat from a transect where *S. olivaceus* were observed. d) Angelina National Forest, *S. consobrinus* are abundant in the Longleaf Pine Forest of deep East Texas, often observed on trunks of trees, stumps and fallen logs.

**Figure 6.**
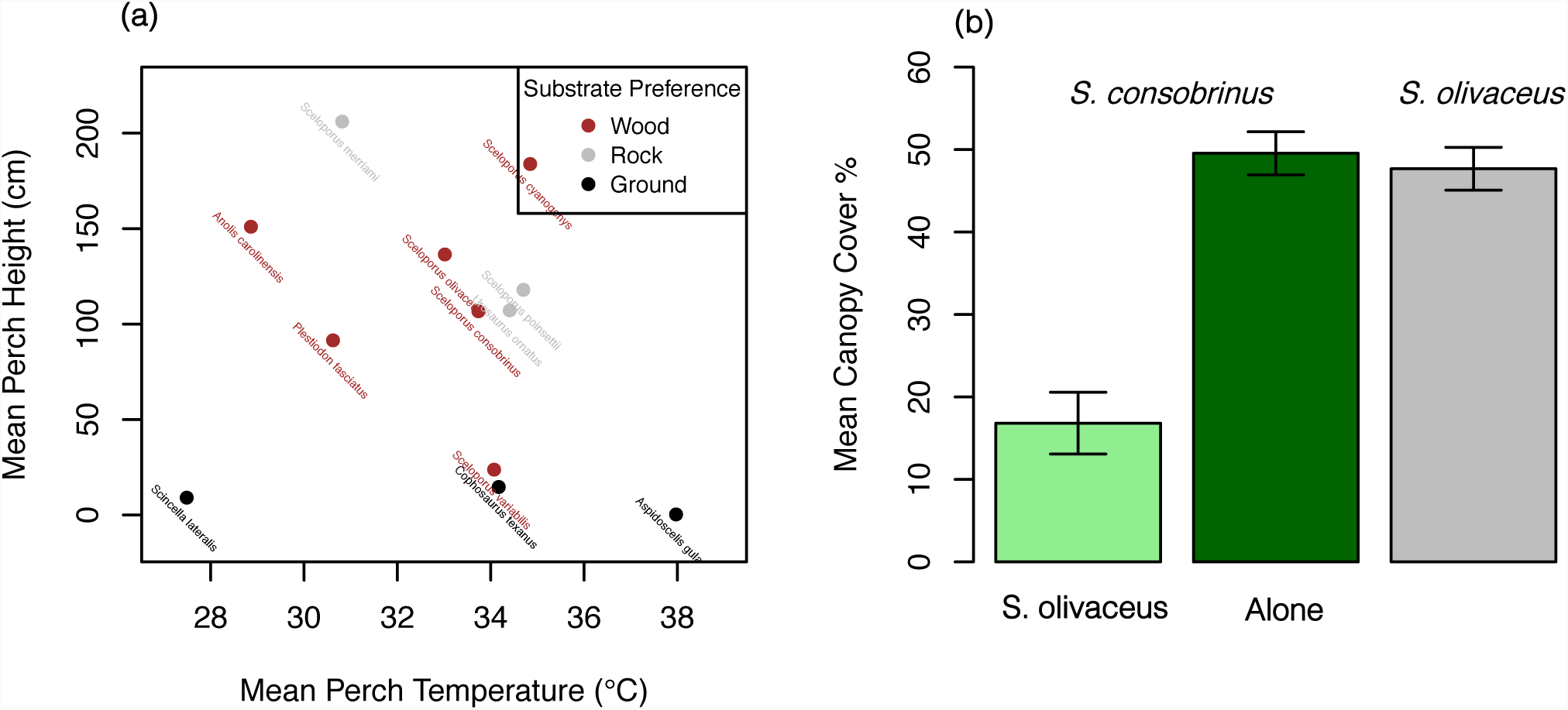
a) Microhabitat use amongst the 12 most common species recorded on transects. Point color represents the substrate which they are most frequently found on, for all species these colors represent greater than 50% of on transect observations). The species mean perch height and temperature values are only from active individuals observed during the day. b). Mean canopy cover of *S. consobrinus* from sites with and without *S. olivaceus*, compared to mean canopy cover for *S. olivaceus*, bars represent standard error.

If competition between *S. consobrinus* and *S. olivaceus* is occurring, we might expect to see character displacement, where *S. consobrinus* alters its habitat use when in the presence of *S. olivaceus*. Perch height is unaffected in this way (p >0.05). But the average canopy cover where *S. consobrinus* is found depends on whether *S. olivaceus* is present in the region. Both *S. olivaceus* across all sites, and *S. consobrinus* at sites where *S. olivaceus* does not occur, occupy transects with similar canopy cover (mean *S. olivaceus* = 47.7%, mean *S. consobrinus* = 49.5%), and the amount of canopy covered used by these species is statistically indistinguishable (p >0.05). But in areas where *S. olivaceus* is present, the two species differ in their habitat use (p-value <0.001), with *S. consobrinus* occurring on much more open transects (16.8 % canopy cover; Figure 6b). As a result, in areas where both species occur, *S. olivaceus* uses more closed canopy environments, while *S. consobrinus* appears to be pushed to open habitats.

At the local level then, there is a signature of potential competition leading to habitat displacement. Is there a similar signature of competition in raw abundances? Overall, *S. consobrinus* was observed on 40 transects, whereas *S. olivaceus* was observed on 58. Frequency of *S. consobrinus* occurrence was lower in regions possessing *S. olivaceus* (22/144, 15%), compared with transects outside of the range of *S. olivaceus* (18/32, 56%). Co-occurrence between *S. olivaceus* and *S. consobrinus* was only observed on 2 of 176 transects, much less than expected by chance. If competition were responsible for *S. consobrinus*’ declines through time, then we would expect present day transects with *S. olivaceus*, to have lower than expected *S. consobrinus* abundance than expected based on their climate suitability for *S. consobrinus*, their canopy cover, and their degree of urbanization. Indeed, abundance of *S. consobrinus* was significantly lower on transects where *S. olivaceus* was present (p-value = 0.024, model R^2^ = 0.57). Urbanization at landscape scales (within 2km of the transect) also negatively impacted *S. consobrinus* (p-value = 0.016). However, corresponding to results presented above, the predicted climate suitability from Maxent models did not have a significant impact on abundance of *S. consobrinus* when all other variables are accounted for (p-value >0.05). Finally, canopy cover alone did not have a significant influence over abundance of *S. consobrinus* (p-value >0.05), *S. consobrinus* was found on transects ranging from 0 to 89.3% canopy cover.

### Competition Trials

Patterns of co-occurrence, when controlling for the environment, can flag potential cases of negative species interactions, but the nature of these interactions are tentative. Specifically, we hypothesize that competition between *S. consobrinus* and *S. olivaceus* is asymmetric, such that historical increases in *S. olivaceus* could cause the declines in *S. consobrinus* (rather than vice versa). We staged competition trials to confirm this asymmetry and establish if *S. olivaceus* is indeed the dominant competitor.

When placed together in the same enclosure *S. olivaceus* initiated more than twice as many antagonistic interactions by doing push-ups, head bobbing, or approaching *S. consobrinus* than vice versa (mean number of interactions initiated: *S. olivaceus* = 3.8, *S. consobrinus* = 1.75, Figure 7a). When responding to these interactions *S. consobrinus* was much more likely to retreat and hide, rather than either not responding, or responding with a counter-interaction (average response scores: *S. olivaceus* mean = 0.15, *S. consobrinus* = −1.1, p-value =0.001, Figure 7b). The negative value for *S. consobrinus* indicates that retreat is frequent, whereas the slightly positive value for *S. olivaceus* is driven largely by their tendency to not respond to antagonistic displays from *S. consobrinus*.

**Figure 7.**
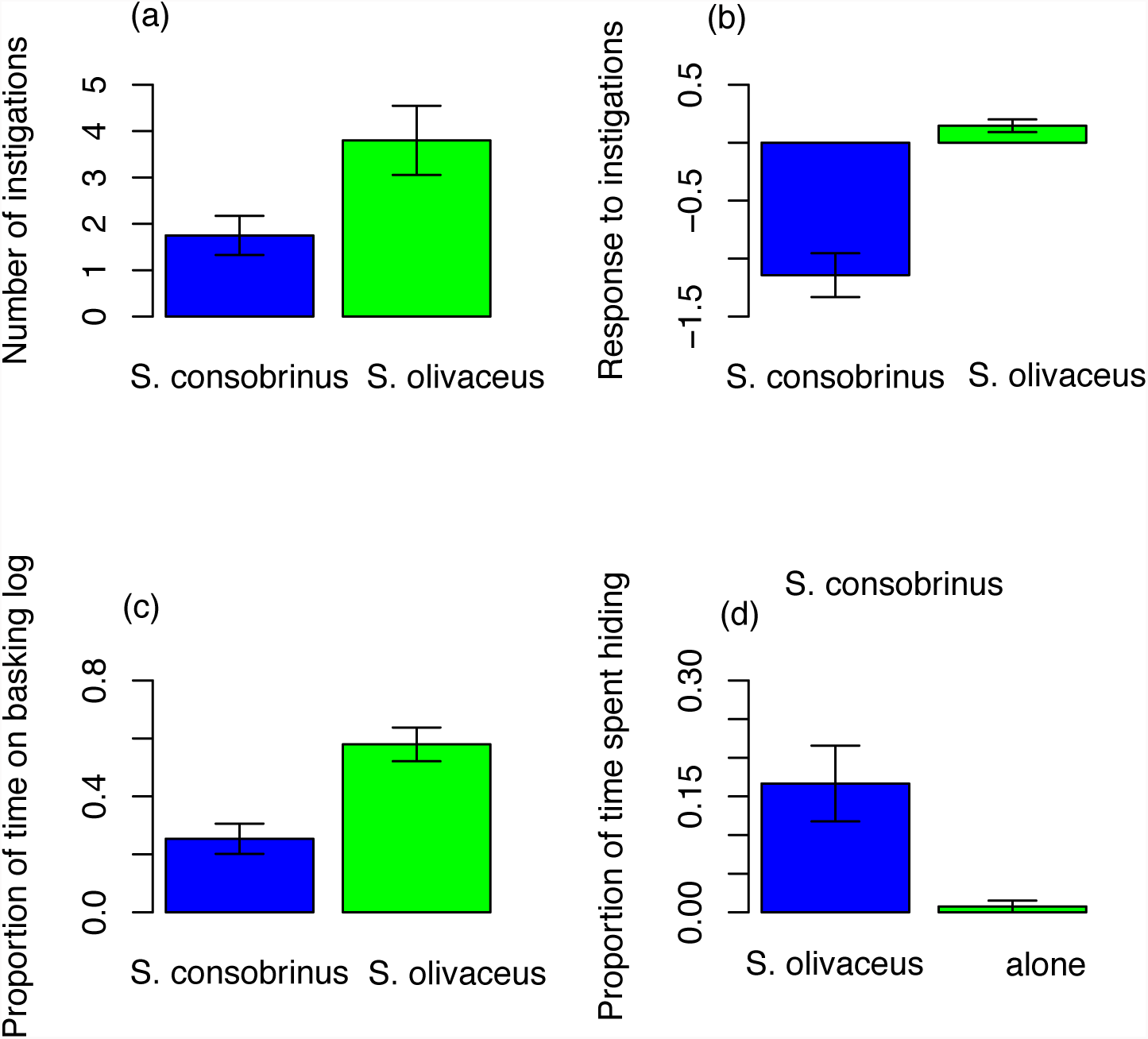
a) Mean number of instigations initiated by each species during behavioral trials, bars represent standard error. b) Mean response to interactions by each species. c) Proportion of time on the basking log during trials. d) Proportion of time spent hiding by *S. consobrinus* in trials with *S. olivaceus* compared to when *S. consobrinus* are alone.

Beyond the types of interactions engaged in, competition trials also revealed that *S. olivaceus* dominated the single available basking location when in the presence of *S. consobrinus* (mean proportion of time on basking log for *S. olivaceus* = 0.58, *S. consobrinus* = 0.25, p-value= <0.001, Figure 7c). In contrast, the amount of time that *S. consobrinus* spent hiding (either under cover objects or by burying itself) increased greatly when *S. olivaceus* was present compared to when *S. consobrinus* was alone (mean proportion of time spent hiding alone = 0.008, with *S. olivaceus* present = 0.17, p-value= 0.004, Figure 7d).

## Discussion

We uncovered a dramatic decline of *S. consobrinus* over the last 100 years (Figure 2a,c) and evaluated major hypotheses to explain this decline. Land use change, namely urban development, negatively impacts *S. consobrinus*, leading to its absence in heavily urbanized areas. However, declines of *S. consobrinus* are not confined to urban areas, with reductions in occurrence taking place in areas far from cities (Figure 3). In such instances, multifaceted evidence across scales points towards interspecific competition with *S. olivaceus* playing a role in excluding *S. consobrinus* from otherwise suitable habitat (Figure 2b). The strength of competition appears to be strong enough to override the measurable influence of climate: predicted occurrence of *S. consobrinus* is equivalent in areas without *S. olivaceus* but with low climate suitability, and areas with extremely high climate suitability which have *S. olivaceus* (Figure 2b). The negative association with *S. olivaceus* likely drives trends of decreasing *S. consobrinus* in natural areas, with co-occurrence with *S. olivaceus* at the pseudo-site level becoming increasingly rare with time (Figure 2c, Figure 3). The conclusions stemming from these broad scale historical records were borne out in transect surveys assessing differences in lizard communities across Texas (Figure 5). Field observations demonstrated a high level of overlap in habitat use between the two species, as *S. olivaceus* uses more similar microhabitat to *S. consobrinus* than any of the other regional species (Figure 6a). In the absence of *S. olivaceus*, the two species are found in habitats with similar canopy cover, but in regions where *S. olivaceus* occurs, *S. consobrinus* occupies far more open areas, effectively partitioning the local habitat (Figure 6b). At the transect level *S. consobrinus* abundance is lower on transects possessing *S. olivaceus* compared with those where it is absent. Despite having similar habitat requirements and the range of *S. consobrinus* subsuming nearly all of *S. olivaceus*’s Texas range, the two species only co-occurred on 2 of 176 transects. Finally, experimental competition trials revealed *S. olivaceus* as a superior competitor to *S. consobrinus*. *Sceloporus olivaceus* were more aggressive, initiating more interactions, and were almost exclusively the winner of interactions (Figure 7a,b). Corresponding to data from transects, we find that the presence of *S. olivaceus* lead *S. consobrinus* to shift their habitat usage, spending over 15% of the time hiding, compared to less than 1% when alone (Figure 7d). These results when taken together suggest that *S. olivaceus* is competitively dominant, capable of excluding *S. consobrinus* via behavioral interference from areas it would otherwise occupy, and that the frequency of such competitive exclusion has been increasing over the last century, to the detriment of *S. consobrinus* populations.

Present day occurrence of *S. consobrinus* is poorly explained by climate alone (Table 1), suggesting that other factors are playing a disproportionately large role in determining their range. Habitat modification, specifically urbanization, is in part responsible for excluding them from otherwise suitable areas (Table 1), and while *S. consobrinus* are largely absent in cities other lizards such as those in the genus *Anolis* can often tolerate urbanization well (Winchell *et al*., 2020). Declines associated with urbanization may stem from many different factors (Shochat *et al*., 2006), as conversion from natural habitats to urban ones can increase temperatures, thereby pushing species past their thermal limits, reduce resource availability, alter availability of microhabitat, and alter species interactions (Shochat *et al*., 2010). However, absences in areas unimpacted by urbanization points to other possibilities. Climate change is often cited as a potential cause of population declines in undisturbed areas (Lister & Garcia, 2018). We found that *S. consobrinus* occurrence was less likely in areas that have experienced warming or had increased in precipitation over the last century (Table 1). Yet, mechanistically, warming seems an unlikely cause, as much of the area in which *S. consobrinus* is absent today have not experienced a great amount of warming. Indeed, once presence of *S. olivaceus* is taken into account, the effect of climate change is no longer significant (Table 1). Our results conform to findings from previous studies have not predicted a decline based off climate alone for *S. consobrinus* (Buckley, 2008). If warming or precipitation changes are playing a role in the current distribution of *S. consobrinus* it may be indirect, in the form of altering biotic factors influencing *S. consobrinus*, such as the strength of competition, reducing prey availability, or altering habitat structure.

Ultimately, the presence of *S. olivaceus* better explains *S. consobrinus* occurrence than any other variable we assessed (Table 1). These results were mirrored at a fine scale by our transect data, showing decreased abundance of *S. consobrinus* on transects containing *S. olivaceus*. Previous studies have demonstrated that competition can lead to exclusion from otherwise suitable areas. For example, urban tolerant bird species are less likely to breed in cities if a superior competitor is present (Martin & Bonier, 2018). Meanwhile species interactions are responsible for species failing to maximize their abundance in accordance with measures of climate suitability (Braz *et al*., 2020). Certainly, other species interactions may be important for *S. consobrinus* beyond that of *S. olivaceus*. *Anolis carolinensis* presence was also negatively associated with *S. consobrinus* based off occurrence record analysis, but at the transect level the two species were not significantly associated with each other, suggesting that caution should be used in scenarios of inferring species interactions from spatially and temporally coarse occurrence data. Signals detected at coarse scales may simply indicate preferences for alternative environments not included in the analysis. For example, *S. consobrinus* and *A. carolinensis* have greatly different preferences in body temperature, and *A. carolinensis* is more heavily arboreal (Figure 6a), and prefers different habitat types, specifically occupying areas with more closed canopy than *S. consobrinus*. Additionally, *Scincella lateralis*, a small and terrestrial lizard, was positively associated with *S. consobrinus* in the occurrence record analysis, yet no relationship was found on transects. Attempts to infer species interactions based off occurrence data must be interpreted with caution (Zurell *et al*., 2018), and ideally collaborated both through time, and at fine spatial scales at which species interactions actually occur.

Competitive interactions have been suggested to be more likely between closely related species (Darwin, 1859), particularly as closely related species tend to have high levels of niche overlap, which itself promotes the likelihood of competition. Our species pair is extremely closely related, possibly capable of hybridization (Smith *et al*., 1991), and have high levels of niche overlap, as both species were found primarily on woody substrates (tree trunks, logs, branches) between 1-1.5m high and 33-34 °C (Figure 6). Behavioral trials between *S. consobrinus* and *S. olivaceus* revealed strong competitive interference, with the much larger *S. olivaceus* clearly dominant (Figure 7). Actual attacks rather than merely aggressive displays were rare. In only one instance did *S. olivaceus* engage physically with *S. consobrinus*, an interaction which resulted in *S. consobrinus* autotomizing its tail. Fear of such consequences, even when attacks are extremely uncommon, may lead to behavioral changes in *S. consobrinus*. Body size is often key in determining competitive hierarchies (Munday *et al*., 2001; Price & Sheilds, 2002), and *S. olivaceus*’ dominance is likely attributable to is larger size. In nature, the risk of predation between larger and smaller lizards (even if actual predation is rare) can lead to behavioral avoidance between lizard species with substantial population and community level consequences (Pringle *et al*., 2019). Such avoidance frequently takes the form of shifts in habitat use (Culbertson & Herrmann, 2019). While shifting habitat use may allow for coexistence of species, such shifts can also have negative consequences, such as altering thermoregulation success or foraging ability. Further, it may not always be possible to partition habitat, and in such instances exclusion of the inferior competitor may occur (Pringle *et al*., 2019). While our data suggests behavioral interference is a factor in excluding *S. consobrinus* from otherwise suitable areas, it is possible that exploitative competition also plays a role. Both species are primarily insectivorous, and likely share prey. However, our study does not assess the influence of prey availability on the rate of co-existence between the two species, nor are we aware of evidence to suggest that populations of these lizards are prey limited.

Interestingly, co-occurrence between *S. olivaceus* on *S. consobrinus* was historically much more common than it presently is. Urbanization seems to contribute to this trend, as in more urban areas *S. olivaceus* is favored while *S. consobrinus* is disfavored. But *S. consobrinus* declines are also occurring in areas far from urban centers. The decline of *S. consobrinus* was first noted in 1991 in an unpublished dissertation from the Welder Wildlife Refuge in Southern Texas (Mora, 1991). This study found that between 1959 and 1988 *S. consobrinus* went from being the most common lizard found in all habitat types on the refuge, accounting for 17% percent of all lizard observations, to only one individual being observed over a two-year period (out of 1,706 lizards observed, i.e. 0.059% of observations). During this time *S. olivaceus* increased greatly, becoming the most widespread lizard in the refuge, and accounting for 38% of all lizard observations in 1988. Other *S. consobrinus* declines are not well documented, however some North Texas populations of *S. consobrinus*, which were extant throughout 1975-2000, (Jones & Ferguson, 1980; Ryberg *et al*., 2005) are now absent (personal observation). In central Texas hill country, a population of *S. consobrinus* which was abundant until 1990 has since been lost, even in the absence of land-use change (David G. Barker personal communication). In all three of these regions *S. olivaceus* remains (or has become) abundant.

While we propose that increasing competition from *S. olivaceus* is largely behind these declines, the exact mechanism behind the increasing competition is unclear. Three potential hypotheses may explain these patterns: (1) Habitat modification across the landscape has reduced habitat heterogeneity, and in a less patchy environment the superior competitor is able to dominate; (2) Decreases in resource availability may cause competition to matter more in determining coexistence; (3) Increasing abundance of *S. olivaceus* has resulted in heightened total competitive pressure on *S. consobrinus*.

Habitat heterogeneity is crucial for the coexistence between competing species, allowing them to partition themselves and avoid negative consequences of competition (Pianka, 1967; Ben-Hur & Kadmon, 2020). In the absence of competitor free space, species are susceptible to competitive exclusion. This may be the case between *S. consobrinus* and *S. olivaceus*. While they have similar habitat use, *S. olivaceus* is more of a habitat specialist than *S. consobrinus*. *S. olivaceus* prefers wooded or forested areas with mature oak, hackberry and mesquite trees (Blair, 1960). They are rarely encountered in areas lacking trees. In the drier western portions of its range *S. olivaceus* is patchily distributed across the landscape, largely confined to forested areas near water bodies (Milstead, 1950). *Sceloporus consobrinus* is more of a generalist and exhibits incredible variation in habitat use throughout its range (Leache & Reeder, 2002), occupying forested habitat when *S. olivaceus* is absent, but capable of using open habitats as well. *Sceloporus olivaceus’s* reliance on trees may explain why climate suitability for *S. olivaceus* is such a strong predictor of *S. consobrinus* absence. Areas where climate suitability is high for *S. olivaceus* may also be regions where *S. olivaceus* is more widespread across the landscape, and thus able to entirely exclude *S. consobrinus* from the region, in comparison to regions where it has a more patchy local distribution. This possibility is consistent with our finding that co-occurrence between the two species is especially unlikely in wetter areas, as compared to dry areas (Figure 4), as wet regions usually are more heavily forested and suitable for *S. olivaceus*. This finding makes sense in light of the decline of *S. consobrinus* at Welder Wildlife Refuge, which occurred over a time with increasing precipitation, and likewise increased density of vegetation across the landscape (Mora 1991).

Beyond habitat heterogeneity, declining resource availability may be dictating the strength of competition between *S. consobrinus* and *S. olivaceus*. In times of low resource availability and environmental stress, such as drought, *S. olivaceus* is able to persist, while other lizards such as *Anolis carolinensis* may be locally extirpated (Blair, 1957). Enhancing *S. olivaceus* ability to deal with fluctuation in resource availability is their larger size which may be key in their ability to switch prey items, even consuming small vertebrate prey such as rough green snakes (Blair, 1960) and *Hemidactylus turcicus* (Personal Observation). Although not well documented, others have suggested that *S. olivaceus* may consume other lizard species (Lawrence LC Jones & Lovich, 2009), and juvenile *S. consobrinus* are small enough to be eaten by large *S. olivaceus*. As such intra-guild predation may occur during periods of low resources, especially impacting *S. consobrinus* given that they have the same general habitat preferences.

Finally, *Sceloporus olivaceus* appears to be increasing in relative abundance over the last 100 years based off occurrence data (Figure 2a), and areas resurveyed over that time period also show large increases of *S. olivaceus* abundance (e.g. Mora, 1991). Increases of *S. olivaceus* may explain decreased rates of co-occurrence with *S. consobrinus* since the net impact of competition is the product of per capita competitive effects and the total number of competitors. When a superior competitor is at higher density, niche breadth of inferior species is more strongly reduced (Tarjuelo *et al*., 2017). Likewise, foraging may be possible when competitors are at low density, but success declines as density increases (Hasegawa, 2016). The mechanism behind the increase in abundance of *S. olivaceus* is ultimately unknown, but may be related to changing precipitation or land use patterns.

## Conclusions

Here we find evidence that *Sceloporus consobrinus* has declined over the last 100 years, and that asymmetric interference competition with *S. olivaceus* is plausibly responsible. These declines lead to a reduction in the distribution of *S. consobrinus*, with many parts of central Texas and south Texas no longer harboring *S. consobrinus* populations. We demonstrate that broad-scale associations between species occurrence patterns from historical records correspond to fine scale ecological data from transect surveys, which also match individual-level behavioral responses recorded from staged competition trials. Across all these scales the data support *S. olivaceus* as playing a role in limiting the distribution and abundance of *S. consobrinus.* We offer insight into the potential mechanisms responsible for an increase in their competition, and suggest that future studies focus on identifying the mechanisms by which species interactions are changing. The role of competition in setting range limits is widely studied, often in the context of invasive species or maintaining boundaries between closely related parapatric species. Our study suggests that the strength of competition can fluctuate through time, and increasing competition between native species with large areas of range overlap, can lead to range loss for the inferior competitor—even in their range interior.

## Acknowledgements

We thank the Nature Conservancy for allowing us to work in exceptionally high quality habitat. We are grateful to Chandler Davis, Dalton Lawing, Dan Nicholson, and Greg Pandelis, who all assisted with fieldwork. We thank Texas Ecolabs for financial support of this work, as well as the private land owners who collaborated through Texas Ecolabs and granted us access to their property. This work was carried out under UTA IACUC number A2019.0007, Texas Parks and Wildlife permit #SPR-0814-159 and #SPR-0323-25, as well as Fish and Wildlife Special Use Permit #75742. The Frishkoff lab was supported by NSF Award #2055486.

