## Supplemental Figures for "Competition and the century-long decline of a once common lizard, *Sceloporus consobrinus*"

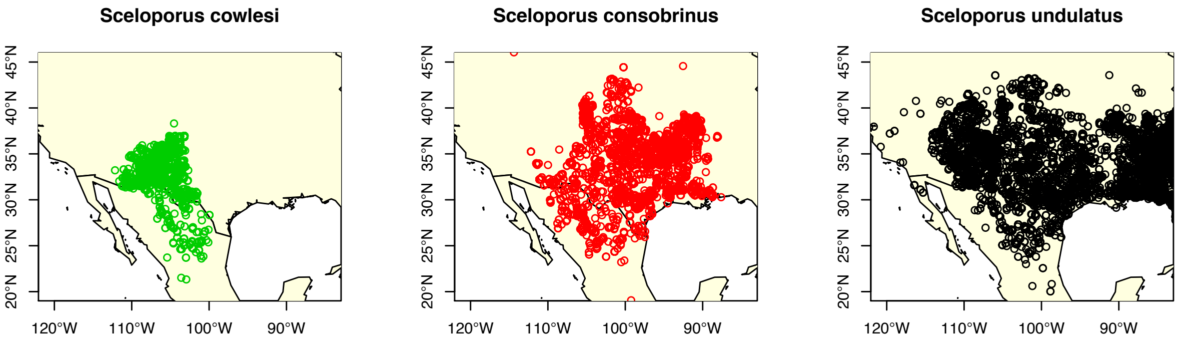


**Figure S1** Occurrences of the *S. undulatus* group on gbif, before data was cleaned and taxonomy updated.

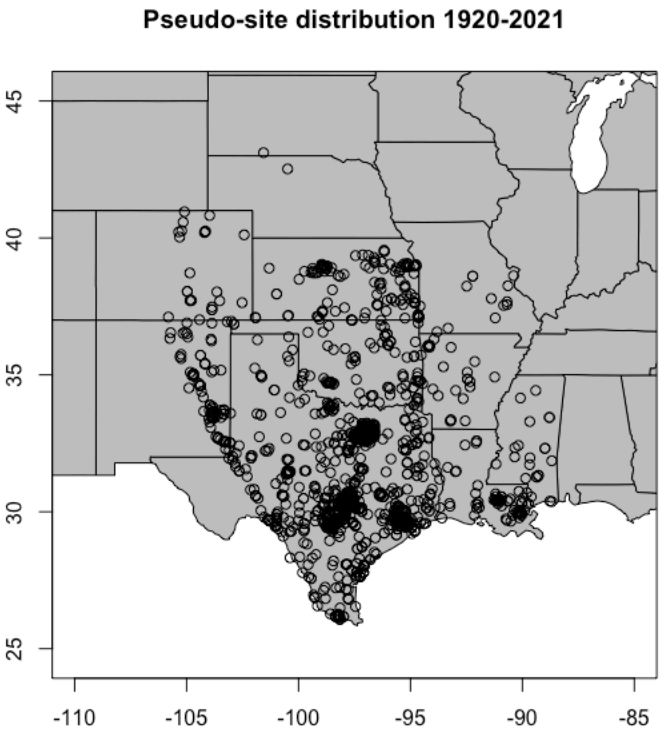


**Figure S2.** Pseudo site distribution at 2km sites, 1340 total records between 1920-2021 with a minimum of 5 observations per location.

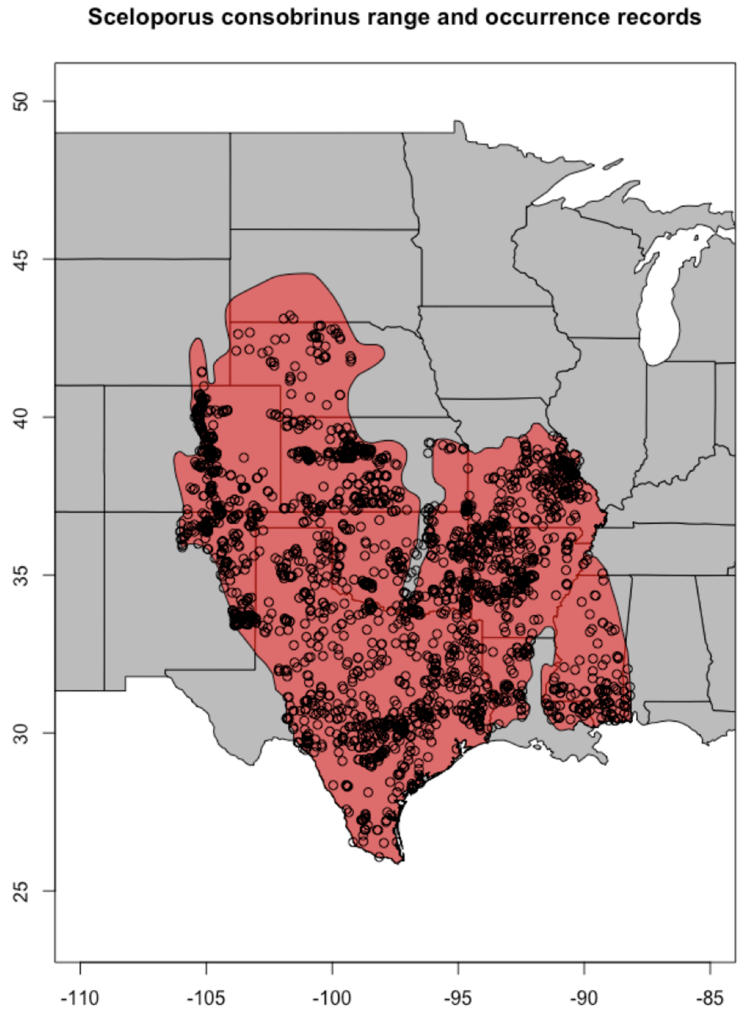


**Figure S3** Cleaned occurrence records, post thinning, used for Maxent models for *Sceloporus consobrinus*. 2527 records

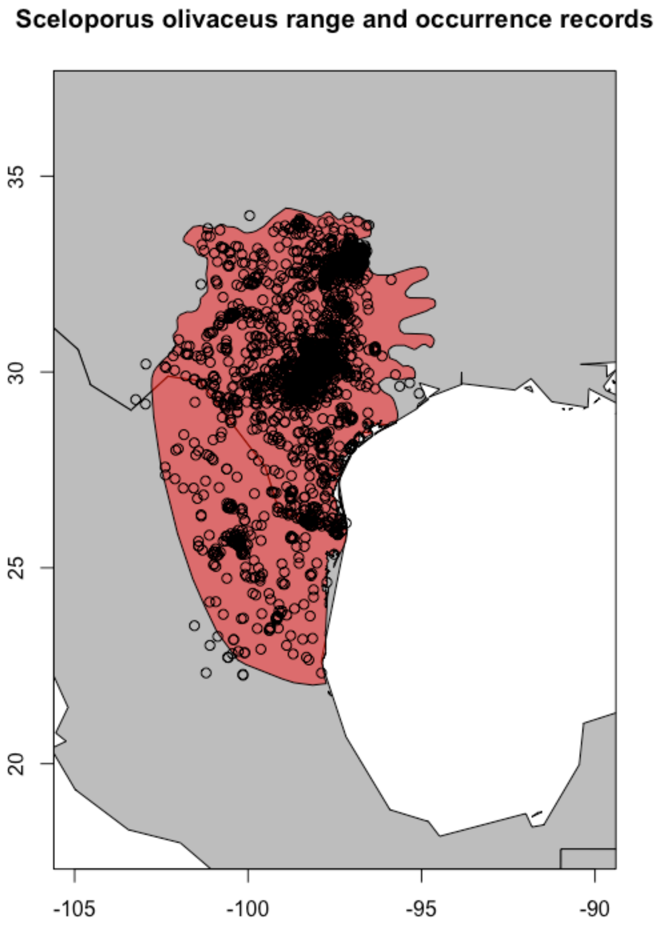


**Figure S4** Cleaned occurrence records, post thinning, used for Maxent models for *Sceloporus olivaceus*. 2817 records.

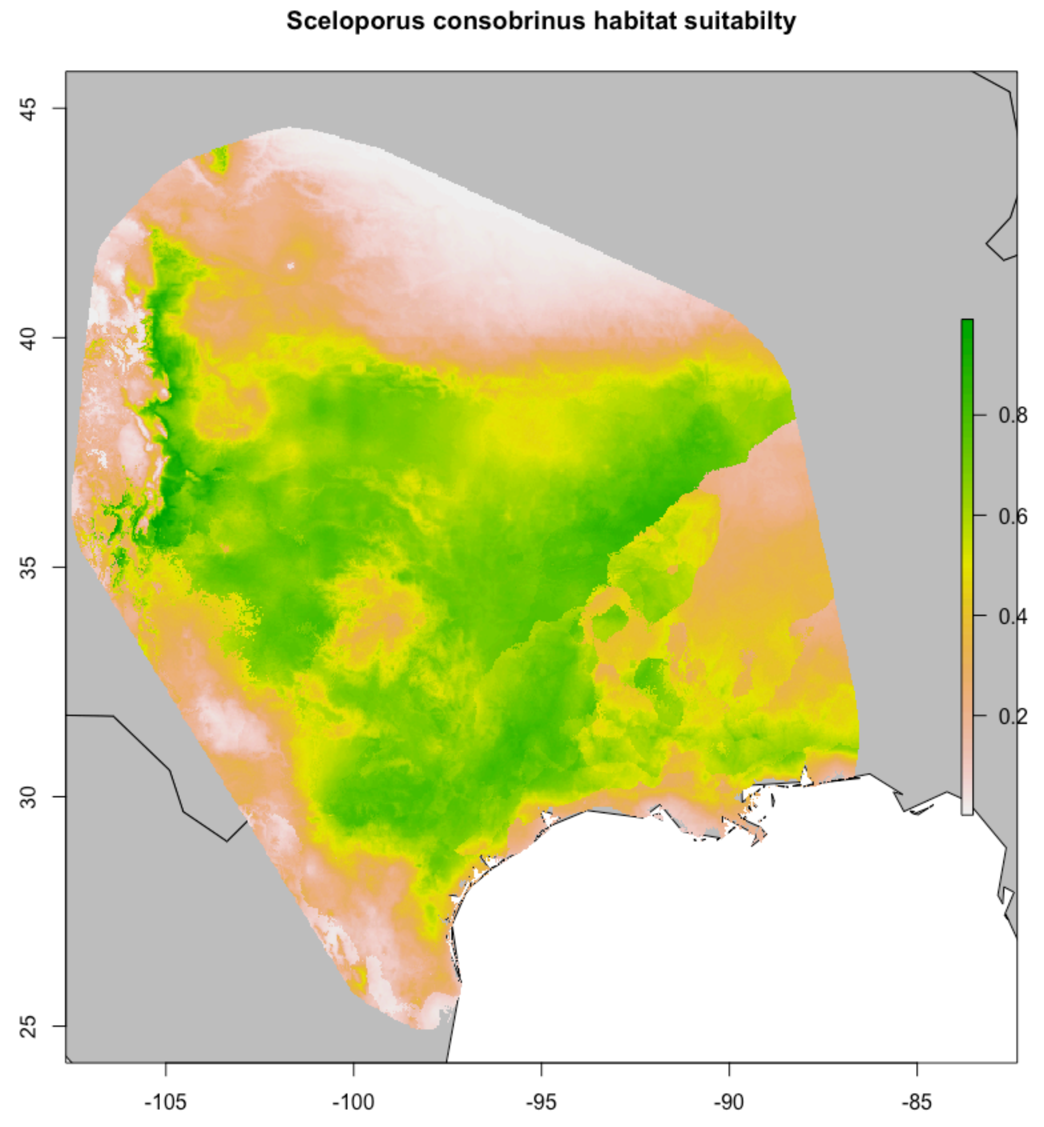


**Figure S5.** Maxent predicted climate suitability for *Sceloporus consobrinus*, AUC value=0.749.

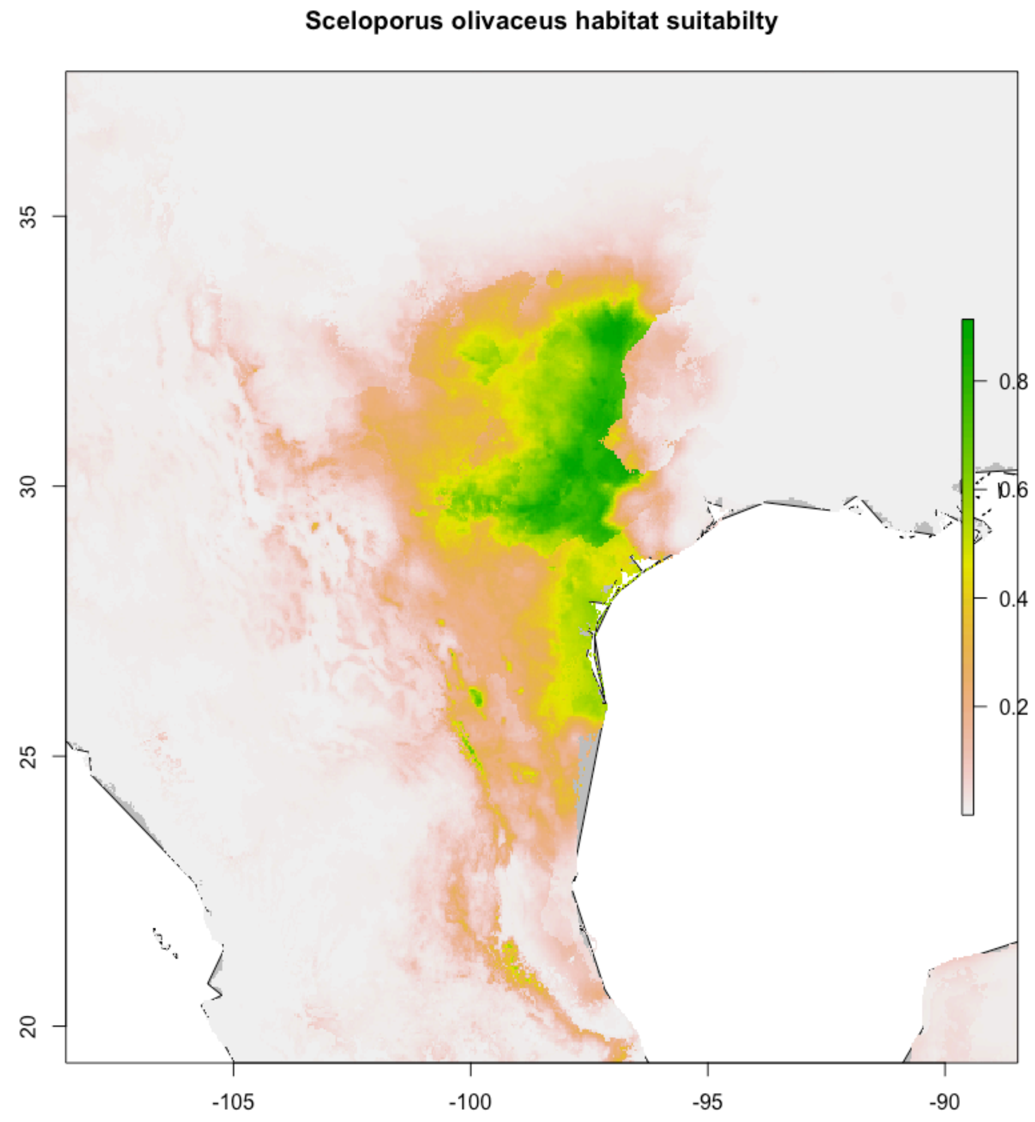


**Figure S6.** Maxent predicted climate suitability for *Sceloporus olivaceus*, AUC value=0.909.

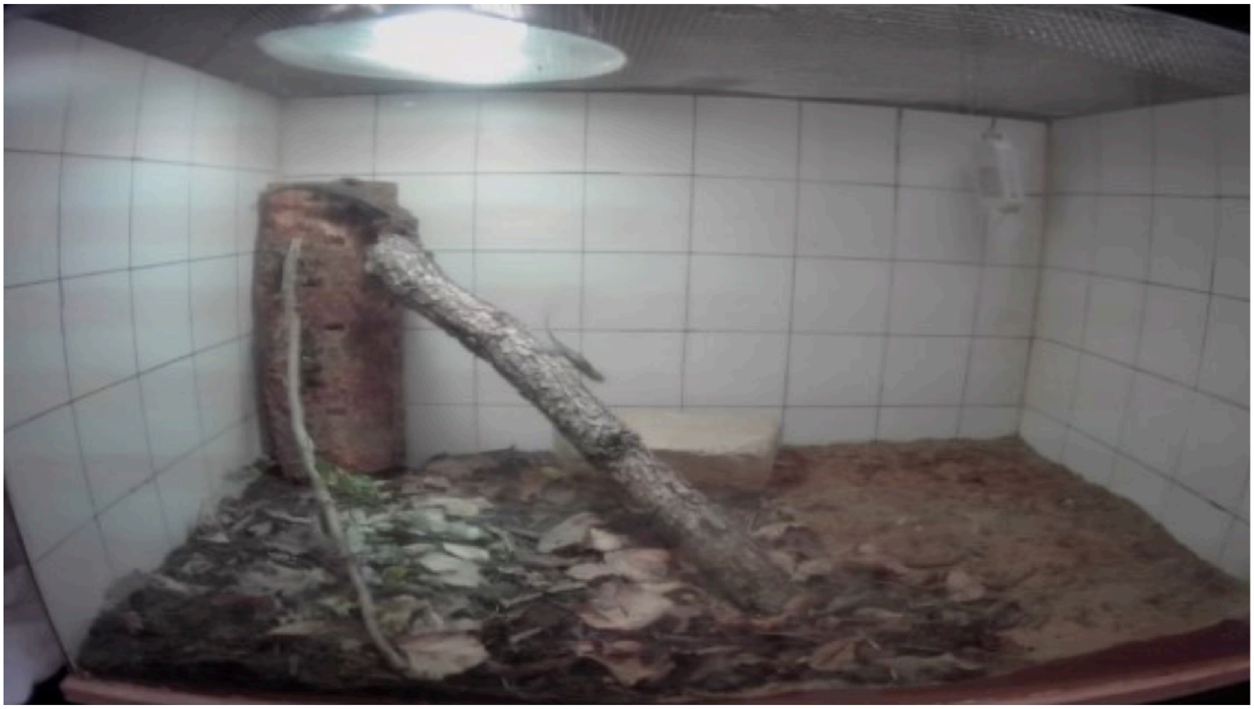

**Figure S7.** Screenshot from a competition trial to illustrate enclosure design. Note large *Sceloporus olivaceus* on top of basking perch in upper left corner, and *Sceloporus consobrinus* at middle of tilted branch in the center of the screen, retreating from a recent behavioral interaction.
